# Stairway Plot v3: Nonparametric Demographic Inference from the Site Frequency Spectrum at Biobank Scale

**DOI:** 10.64898/2026.09.16.751895

**Authors:** Xiaoming Liu

## Abstract

Inferring the history of effective population size (*N*_*e*_) from genomic data is fundamental to population genetics, but model-flexible methods have not kept pace with biobank-scale samples. We present Stairway Plot v3, which infers a nonparametric *N*_*e*_(*t*) trajectory with bootstrap confidence intervals directly from the site frequency spectrum (SFS). By recasting the optimization as a convex problem and combining inferences across multiple sub-sampled projections, the method scales to 10^5^ haploid sequences (*n*_seq_) on commodity CPU hardware without a GPU. Larger samples sharpen the resolution of recent demography, and we apply the method to five human populations including 54,302 Japanese individuals, to our knowledge the largest sample used for nonparametric SFS-based *N*_*e*_(*t*) inference.

---

The effective population size through time, *N*_*e*_(*t*), records the demographic history of a population—its expansions, bottlenecks and separations—and is among the most widely estimated quantities in population genetics. Genomic datasets have grown to hundreds of thousands of sequenced individuals, and these large samples carry information about very recent demography that small samples cannot, because recent population-size changes are written into rare, low-frequency variants that only large samples observe in abundance. Yet the model-flexible methods most often used to estimate *N*_*e*_(*t*) without pre-specifying a demographic model were designed for small samples: the sequentially Markovian coalescent methods PSMC^1^ and MSMC^2^ analyze 1–4 diploid genomes, SMC++^3^ incorporates more samples but still tracks only a few distinguished lineages, Relate^4^ builds genome-wide genealogies for all haplotypes, PHLASH^5^ extends the PSMC framework with GPU-accelerated Bayesian sampling. Stairway Plot^6,7^ takes a different route, inferring a piecewise-constant *N*_*e*_(*t*) (also known as constant-in-state model) directly from the SFS—a summary statistic whose size depends only on the sample, not on the length of the genome, and whose computation is trivially parallelizable. FitCoal^8^ also fits a constant-in-state demographic model to the expected SFS with AIC-based model selection. None of these was built to exploit, or even to ingest, biobank-scale samples.

Here we present Stairway Plot v3, which makes this constant-in-state modeling of SFS practical at biobank scale through two changes. First, we replace the stochastic differential-evolution optimizer of previous versions with a convex interior-point solver. The expected SFS proportions are linear in the scaled mutation-rate parameters *θ*_*j*_ = 4*N*_*e,j*_*μ*, so the full multinomial log-likelihood is concave in ***θ*** (Methods). For a fixed set of epoch groups, a Newton-based barrier method therefore reaches the global optimum in 10–20 iterations rather than the thousands of black-box evaluations differential evolution requires; model selection over breakpoint placements remains heuristic and is explored by replicated randomized searches.

Second, because estimating one parameter per coalescent interval is intractable for a sample of 10^5^, we infer *N*_*e*_(*t*) at several smaller down-projected sample sizes (projections) and combine them with a sigmoid weighting that assigns recent epochs to large projections and ancient epochs to small ones, yielding a single trajectory with bootstrap confidence intervals (Methods). Together these keep inference memory-light and GPU-free: on a single CPU workstation it completes in minutes at *n*_seq_ ≤ 10^3^ and in about five hours at *n*_seq_ = 10^4^ (8 threads). Because the bootstrap replicates are independent, the computation parallelizes near-linearly across CPU cores or cluster nodes, making *n*_seq_ = 10^5^ feasible on a single workstation (~67 h with 156 GB RAM) and routine on a cluster.

To benchmark the method we simulated three demographic models—a European bottleneck-and-expansion (CEU), an African expansion (YRI) and an oscillating “zigzag” history—with msprime^9^, and compared Stairway Plot v3 against its predecessor Stairway Plot 2^7^ and three recent methods, FitCoal^8^, PHLASH^5^ and Relate^4^, at sample sizes of *n*_seq_ = 100, 1,000 and 10,000 (Fig. 1a, Table 1; extended to *n*_seq_ = 20,000 in Supplementary Fig. 1). We did not re-compare MSMC2 or SMC++, which were benchmarked against Stairway Plot 2 previously^7^ and do not scale to the larger sample sizes that are the focus here. At *n*_seq_ = 100 the methods performed broadly comparably, with none dominating across models. The advantage of Stairway Plot v3 emerged as samples grew: at *n*_seq_ = 1,000 it gave the lowest error on all three models, roughly halving the error of Stairway Plot 2 on the CEU and zigzag models, while running in ~8 min versus ~55 min for v2’s differential-evolution optimizer and tens of hours for the genealogy- and coalescent-HMM methods. At *n*_seq_ = 10,000, under our benchmark configuration (default parameters, shared 156 GB workstation), only Stairway Plot v3 completed: Stairway Plot 2’s optimizer did not scale, Relate’s genealogy-building exceeded available storage (~1.5 TB of intermediate files), PHLASH stalled before its first sampling iteration on a 12 GB GPU, and FitCoal was not run, as it becomes computationally impractical at this sample size.

**Table 1.** Accuracy (mean squared log error, MSLE; lower is better) and wall-clock time across methods and sample sizes.

| Method | MSLE (CEU / YRI / zigzag) |  |  | Wall time |  |  | Hardware |
| --- | --- | --- | --- | --- | --- | --- | --- |
| | $n=100$ | $n=1,000$ | $n=10,000$ | $n=100$ | $n=1,000$ | $n=10,000$ | |
| SP v3 | 0.11/**0.03**/0.02 | **0.02/0.01/0.01** | **0.01/0.01/0.01** | ~7 min | ~8 min | ~5 h | CPU |
| SP v2 | 0.08/0.03/**0.02** | 0.04/0.01/0.02 | — | ~11 min | ~55 min | — | CPU |
| Relate | **0.06**/0.04/0.04 | 0.06/0.06/0.05 | — | 2–4 h | 38–50 h | — | CPU |
| PHLASH | 0.27/0.06/0.10 | 0.29/0.05/0.05 | — | ~54 min | ~52 min | — | GPU |
| FitCoal | 0.27/0.19/0.13 | 0.17/0.02/0.17 | — | ~10 h | 82–94 h | — | CPU |

MSLE entries are CEU / YRI / zigzag (lower is better), computed on a log-uniform time grid; bold marks the lowest error for each model at each sample size. At $n_{\text{seq}} = 100$ the best method differs by model (Relate on CEU, SP v3 on YRI, SP v2 on zigzag); from $n_{\text{seq}} = 1,000$ onward SP v3 is best on all three. SP v2 = Stairway Plot 2 (v2.1.3, differential-evolution solver). A dash (—) indicates the method did not complete at that sample size with default settings (SP v2: differential-evolution optimizer did not scale; Relate: ~1.5 TB intermediate storage; PHLASH: optimization stalled on a 12 GB GPU; FitCoal: not run). SP v3 at $n_{\text{seq}} = 10,000$ used 8 threads.

**Figure 1.**
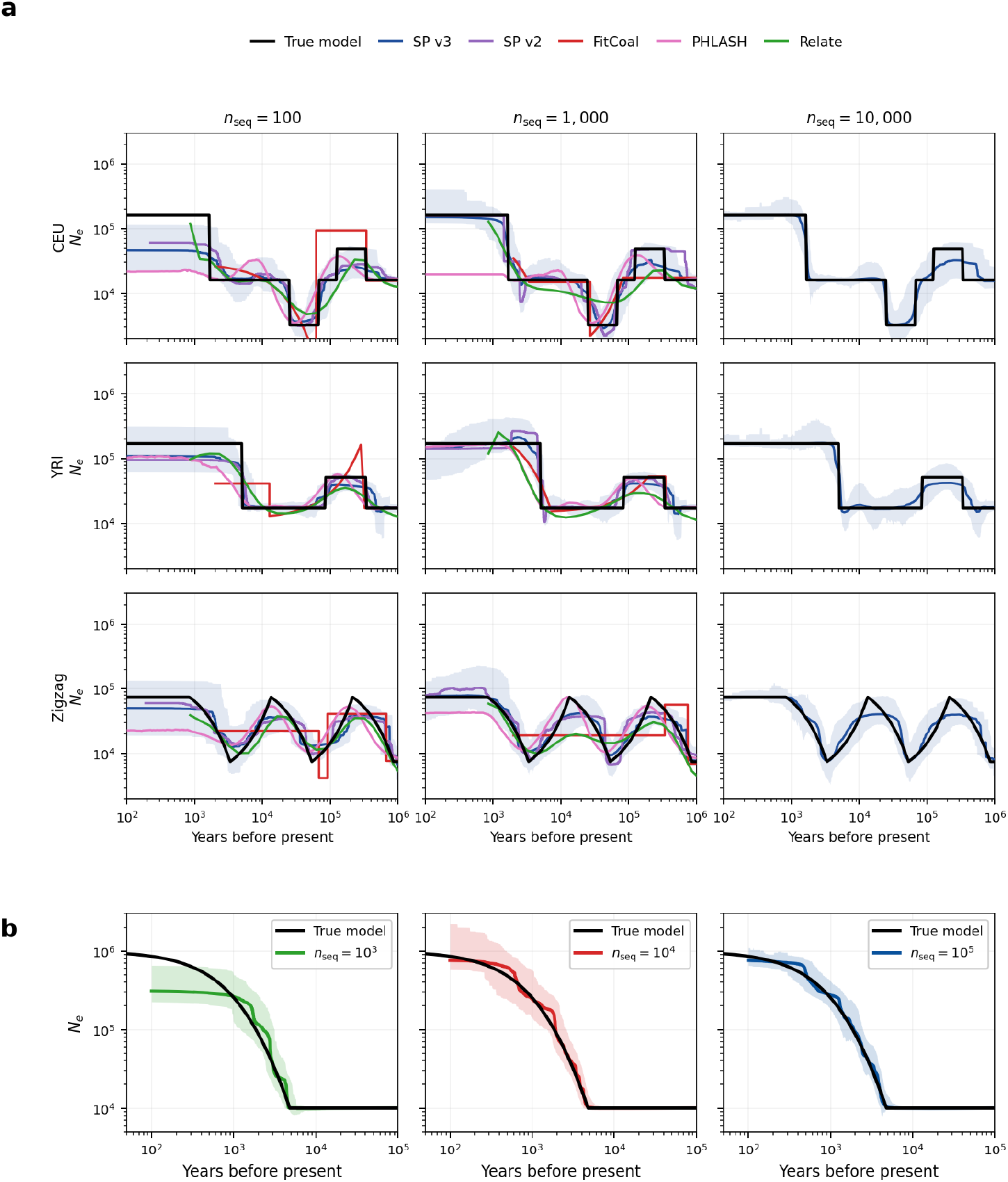
Stairway Plot v3 scales to biobank-sized samples and resolves recent demography. (**a**) Benchmark on three simulated models (rows: CEU, YRI, zigzag) at three sample sizes (columns: *n*_seq_ = 100, 1,000, 10,000). Each panel overlays the true model (black) with Stairway Plot v3 (blue, 95% bootstrap CI shaded), its predecessor Stairway Plot 2 (purple), FitCoal (red), PHLASH (pink) and Relate (green). v3 and v2 are comparable at *n*_seq_ = 100 but v3 is more accurate at 1,000; at *n*_seq_ = 10,000 only Stairway Plot v3 completes—Stairway Plot 2 does not scale, Relate exceeds storage, PHLASH stalls, and FitCoal was not run. (**b**) Accuracy at scale: a super-exponential expansion (*N*_*e*_: 10^4^ → 10^6^ over the last 200 generations) inferred from identical data projected to *n*_seq_ = 10^3^, 10^4^ and 10^5^ (folded SFS, 95% bootstrap CIs); the recovered recent peak approaches the truth as sample size grows.

The benchmark models above involve no strong recent expansion, so their SFS is informative even at modest samples and larger *n* yields little. To test whether biobank-scale samples deliver the recent-time resolution they should, we simulated a super-exponential expansion—a population growing from *N*_*e*_ = 10^4^ to *N*_*e*_ = 10^6^ over the most recent 200 generations—and inferred its history from the *same* simulated data projected to *n*_seq_ = 10^3^, 10^4^ and 10^5^ (Fig. 1b). The recovered recent peak rose sharply from *n*_seq_ = 10^3^ (*N*_*e*_ ≈ 3.1 × 10^5^) to 10^4^ (7.6 × 10^5^) and plateaued at 10^5^ (7.6 × 10^5^), while the confidence interval continued to narrow with increasing sample size. Every projection correctly recovered the ancient *N*_*e*_. Resolution of recent demography—both point estimates and uncertainty—thus continues to improve up to 10^5^ sequences, exactly the regime that existing model-flexible methods cannot reach.

We recommend folded SFSs for real-data analyses because they are invariant to errors in ancestral-allele assignment, to which only the unfolded spectrum is exposed. Supplementary Figs. 1 and 2 compare folded and unfolded inference in simulation, where ancestral alleles are known exactly.

We applied Stairway Plot v3, using the folded SFS, to five human populations spanning three orders of magnitude in sample size (Methods). As a stringent test of recent-time resolution, we compared the inferred histories of three East Asian populations to probe the contested origin of the Yayoi, the major ancestral component of modern Japanese (Fig. 2). Using 54,302 Japanese individuals (54KJPN^10^; *n*_seq_ = 108,604, to our knowledge the largest sample used for nonparametric SFS-based *N*_*e*_(*t*) inference), 4,157 Koreans (Korea4K^11^) and 4,480 southeastern Chinese (WBBC^12^, mostly from Zhejiang and Jiangxi provinces), we found that at ancient times all three trajectories overlap, consistent with shared ancestry, but in the recent period the 95% bands of the southeastern-Chinese and Japanese trajectories separate at several points within the last ~9 kya, whereas the Japanese and Korean trajectories overlap throughout except at the most recent time points. This greater demographic similarity of the Japanese to the Korean than to the southeastern-Chinese population is consistent with greater shared recent history with the Korean reference population, in line with recent ancient-genome studies^13,14^. Non-overlap of pointwise bands is not a formal test of trajectory equality and does not by itself establish a migration route. Such inferences are necessarily indirect—modern populations are proxies for ancestral sources, and a panmictic *N*_*e*_(*t*) model cannot represent admixture or structure—so they are best read as a complement to genetic-structure and ancient-DNA evidence. Two additional cohorts further illustrate the method’s resolution (Fig. 2c,d). The Ashkenazi-Jewish trajectory (gnomAD v4; 1,736 individuals, *n*_seq_ = 3,472) recovers the known medieval founder event^15^ and reveals a putative bottleneck at ~10 kya (*N*_*e*_ ≈ 15,000), which would predate the formation of the Ashkenazi community and likely reflects its ancestral source population. The Finnish trajectory (gnomAD v4; 5,316 individuals, *n*_seq_ = 10,632) recovers the characteristic founding bottleneck at ~3.5 kya (*N*_*e*_ ≈ 14,400)^16^.

**Figure 2.**
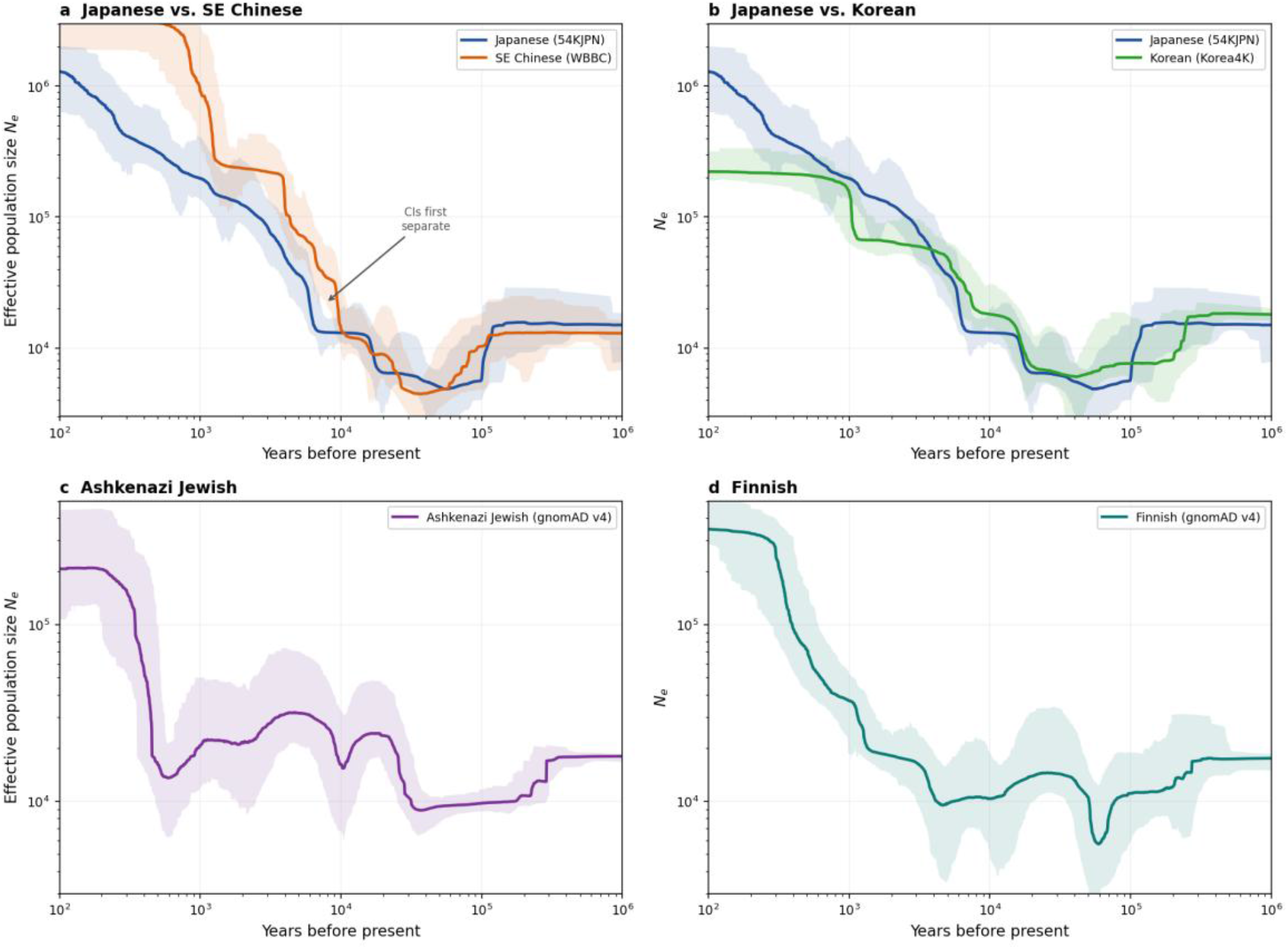
Biobank-scale real-data applications. (**a**) Japanese (54KJPN, *n*_seq_ = 108,604) vs. southeastern Chinese (WBBC): the 95% CIs show a sustained separation around 7–9 kya, consistent with less shared recent history. (**b**) Japanese vs. Korean (Korea4K): the 95% bands overlap through most of history, consistent with greater shared recent history with the Korean reference population. (**c**) Ashkenazi Jewish (gnomAD v4): a putative bottleneck at ~10 kya (*N*_*e*_ ≈ 15,000) is resolved, likely predating the Ashkenazi community and reflecting its ancestral source population, distinct from the medieval founder event visible in the recent period. (**d**) Finnish (gnomAD v4): the characteristic ~3.5 kya bottleneck (*N*_*e*_ ≈ 14,400) is recovered, consistent with the known founder effect of the ancestral Finnish population. All analyses use the folded SFS with 95% bootstrap CIs.

By giving the wider research community a fast, nonparametric and uncertainty-aware estimator of *N*_*e*_(*t*) that runs on commodity hardware at biobank scale, Stairway Plot v3 brings demographic inference into step with the size of modern genomic datasets. The method is distributed as a Python command-line tool wrapping a Java inference engine, with single-command local execution and cluster support (https://github.com/xiaoming-liu/stairway-plot-v3).

## Methods

### The Stairway Plot v3 algorithm (summary)

Throughout, *n*_seq_ denotes the number of haploid sequences (twice the number of diploid individuals). Stairway Plot models *N*_*e*_(*t*) as piecewise-constant with change points at coalescent event times (also known as the constant-in-state model)^17,18,19,20,21^. Each coalescent epoch *i* is the interval during which there are *i* ancestral lineages. To reduce the number of free parameters, contiguous epochs are grouped; all epochs in group *j* share a common effective population size *N*_*e,j*_ and scaled mutation rate *θ*_*j*_ = 4*N*_*e,j*_*μ*. The expected site frequency spectrum (SFS) is linear in ***θ***, *p*_*k*_(***θ***) = *∑*_*j*_ *a*_*kj*_ *θ*_*j*_, where *a*_*kj*_ is the expected branch length at frequency *k* contributed by the coalescent epochs in group *j* (defined in Supplementary Methods)^22^, so the full multinomial SFS log-likelihood, including the monomorphic class, is concave in ***θ*** (see below). Stairway Plot v3 differs from Stairway Plot 2 in five respects, each detailed in Supplementary Methods: (i) a Newton-based interior-point (barrier) solver^23^ that exploits this concavity to reach the global optimum of the fixed-grouping subproblem in 10–20 iterations, replacing the stochastic differential-evolution (DE) optimizer^24^; (ii) candidate breakpoints sampled uniformly in log_10_ coalescent time, with *n*_rand_ = max(20, ⌊5log_2_*n*_seq_⌉), giving even temporal resolution; (iii) multi-projection sigmoid stitching, in which the SFS is projected to a geometric set of smaller sample sizes {20,50, …,200,000} capped at *n*_seq_, inferred independently, and combined as a sigmoid-weighted geometric mean in log-time so that large projections inform recent epochs and small projections inform ancient ones; (iv) a per-replicate SFS bootstrap (multinomial resampling of the frequency-class counts, conditional on the observed number of segregating sites and assuming independent sites, followed by random hypergeometric projection) so that confidence intervals reflect multinomial sampling, projection and fitting variability, but not linkage among sites; and (v) forward–backward–swap model search with smoothness and curvature regularization. Confidence intervals are the 2.5–97.5th (95%) and 12.5–87.5th (75%) percentiles across *B* = 200 replicates, stitched per-percentile.

### Conditioning and the monomorphic class

The software optimizes the full multinomial log-likelihood over all *L* sites, including the monomorphic class:

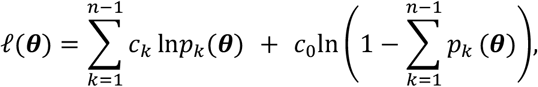

where *c*_*k*_ is the observed count at frequency *k*, 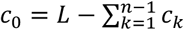 is the number of monomorphic sites, and *p*_*k*_(***θ***) = *∑*_*j*_ *a*_*kj*_ *θ*_*j*_ is the expected proportion of sites at frequency *k*. The input SFS contains only the segregating-site counts *c*_1_, …, *c*_*n*−1_; the monomorphic count *c*_0_ is computed internally from *L*. The constraint *∑*_*k*_ *p*_*k*_ < 1 ensures the monomorphic class retains positive probability. Since *p*_*k*_ is linear in ***θ*** and ln(⋅) is concave, both the segregating-site terms and the monomorphic term *c*_0_ln(1 − *∑p*_*k*_) are concave in ***θ***, so *l* is concave. For the folded SFS, folded-bin probabilities are sums of complementary unfolded-bin probabilities (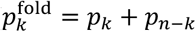 for *k* < n/2), which remain linear in ***θ***; concavity therefore holds for both folded and unfolded likelihoods. For real-data analyses, *L* is the total number of autosomal intergenic sites passing the site mask (see SFS construction below).

### Simulations

Simulations were performed with msprime v1.3^9^ using mutation rate *μ* = 1.2 × 10^−8^ per site per generation^25,26,27^, per-base per-generation recombination rate *r* = 1.2 × 10^−8^, generation time *g* = 24 years^28^, and 10 chromosomes of 30 Mb each (total *L* = 300 Mb), with random seed 42. Three demographic models were used for the benchmark: CEU (European, with an out-of-Africa bottleneck and recent growth), YRI (African, with an ancestral expansion), and the oscillating “zigzag” model of Schiffels & Durbin^2^. Folded SFS were used for all Stairway Plot inferences.

### Accuracy-at-scale (super-exponential) experiment

To test whether biobank-scale samples improve the resolution of recent demography, we simulated a super-exponential expansion: *N*_*e*_ constant at 10^4^ before 200 generations ago, then growing as *d*N/*d*t = *α* N^*β*^ with *β* = 1.2 to *N*_*e*_ = 10^6^ at the present (a 4,800-year growth window), where *α* is determined by the boundary conditions (*N*_*e*_ = 10^4^ at *t* = 200 generations, *N*_*e*_ = 10^6^ at *t* = 0). A single dataset was simulated at *n*_seq_ = 10^5^ (10 chromosomes × 30 Mb; same *μ, r, g*), and the resulting SFS was hypergeometrically projected down to *n*_seq_ = 10^4^ and 10^3^. All three sample sizes therefore derive from identical simulated data, isolating the effect of sample size on temporal resolution.

### Benchmarking against other methods

We compared Stairway Plot v3 against its predecessor **Stairway Plot 2** (v2.1.3, DE solver) and three recent methods. **FitCoal v1.2**^8^ was run on the unfolded SFS (its default and recommended mode) with AIC-based model selection; it does not produce confidence intervals. Stairway Plot v3 used the folded SFS throughout; because ancestral alleles are known exactly in simulation, folding discards information but does not introduce bias, making this a conservative test of v3’s accuracy. **PHLASH**^5^ (v1.0.6) was run on pairs of haplotypes with GPU acceleration (NVIDIA RTX 3060, 12 GB) and produces Bayesian credible intervals. **Relate v1.2.4**^4^ was run with phased haplotypes (10 EM iterations, 10 threads); it does not produce *N*_*e*_(*t*) confidence intervals. All methods used identical mutation rate, generation time and input data. We did not re-compare MSMC2 or SMC++, which were benchmarked against Stairway Plot 2 previously^7^ and do not scale to the larger sample sizes that are the focus here.

At *n*_seq_ = 10,000 (5,000 diploids, 10 chromosomes), three methods did not complete: Relate’s tree-building consumed ~146 GB of intermediate files per chromosome (~1.5 TB total), exceeding available storage; PHLASH’s optimization did not progress past its first sampling iteration after several hours under both single- and dual-GPU configurations; and Stairway Plot 2’s DE optimizer did not scale. FitCoal was not run at this sample size. Wall times were measured on a shared workstation with dual Intel Xeon Silver 4108 CPUs (32 cores total), 156 GB RAM and NVIDIA RTX 3060 (12 GB) GPUs; parallelism differed by method (FitCoal single-threaded; Relate 10 threads; PHLASH one GPU; Stairway Plot v3 multiple CPU worker threads, with the *n*_seq_ = 10,000 run using 8). These methods may perform differently with alternative configurations or hardware; we report results obtained with default parameters. A comparison of Stairway Plot v3 folded and unfolded inference across *n*_seq_ = 200 to 20,000 is shown in Supplementary Figs. 1 and 2.

### Accuracy metric

Accuracy was quantified as the mean squared log_10_ error (MSLE), 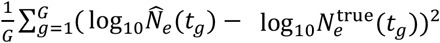, evaluated on a log-uniform time grid of *G* = 500 points (100 to 10^6^ years for CEU/YRI/zigzag; 10 to 10^5^ years for the super-exponential model), with both estimated and true trajectories linearly interpolated in log_10_-log_10_ space. Evaluating on a log-uniform grid—rather than at the native, unevenly spaced estimation points—prevents the metric from being dominated by time regions where a method happens to place many breakpoints. For each method we used its native output: the stitched multi-projection median for Stairway Plot v3, the aggregated final trajectory for Stairway Plot 2, and the point estimate for FitCoal, PHLASH and Relate.

### Real-data applications

Folded SFS were obtained for five populations: **54KJPN**^10^ (54,302 Japanese individuals, *n*_seq_ = 108,604); **Korea4K**^11^ (4,157 Korean individuals, *n*_seq_ = 8,314); **Westlake BioBank for Chinese (WBBC)**^12^ (4,480 southeastern-Chinese individuals, mainly from Zhejiang and Jiangxi provinces, *n*_seq_ = 8,960); and **gnomAD v4** Ashkenazi Jewish (1,736 individuals, *n*_seq_ = 3,472) and Finnish (5,316 individuals, *n*_seq_ = 10,632). All analyses used *μ* = 1.2 × 10^−8^ per site per generation^25,26,27^ and *g* = 24 years^28^. The folded SFS was used throughout because it is invariant to errors in ancestral-allele assignment.

### SFS construction and site filtering

For all five populations, SFSs were constructed from whole-genome sequencing (WGS) allele-frequency databases. Only autosomal chromosomes were included, and only variants with a “PASS” filter status were retained. To restrict the analysis to putatively neutral regions, we defined an intergenic mask by excluding all gene bodies and their flanking 5 kb upstream and downstream regions as annotated in GENCODE v49. The resulting reference intergenic length, *L* = 735,864,277 bp (GRCh38 coordinates, GENCODE v49), was used for all five populations. Cohort-specific callable-region masks were not intersected, so L is a reference intergenic length rather than a per-cohort callable length. Because L enters the likelihood only through the monomorphic count, a proportional error in L shifts the inferred *N*_*e*_ and the time axis by approximately the same factor without changing the shape of the trajectory on logarithmic axes. Related individuals were handled by each database’s own quality-control pipeline. For gnomAD v4, only genome (not exome) data were used.

### SFS imputation

Because the allele-frequency databases report per-site allele counts but may contain missing genotypes, we imputed the SFS using sfs-impute (https://github.com/xiaoming-liu/sfs-impute), which implements the EM algorithm of Liu^29^. This approach iteratively estimates the full SFS from sites with varying levels of missingness, treating the unobserved genotypes as missing data in the EM framework. The imputed folded SFS was used as input to Stairway Plot v3 for all five populations.

## Code and data availability

Stairway Plot v3 (version 3.8.3) is available at https://github.com/xiaoming-liu/stairway-plot-v3 as a Python command-line tool wrapping a Java inference engine (Python 3.8+, Java 8+), supporting single-command local execution and cluster-based parallelization. Real-data SFS were derived from the 54KJPN (https://jmorp.megabank.tohoku.ac.jp/downloads/tommo-54kjpn-20230626-af_snvindelall), Korea4K (https://koreangenome.org/Korea4K_Genomes), WBBC (https://cpbb.cn/resources/GRCh38) and gnomAD v4 (https://gnomad.broadinstitute.org/data) resources.

## AI tools usage statement

Claude Code (Anthropic, Claude Opus 4.6, accessed June 2026) was used as a coding assistant during development of the ensemble and preparation of benchmark scripts. All AI-generated code was reviewed, tested, and modified by the author, who takes full responsibility for its correctness and originality. The manuscript text was drafted by the author with AI assistance. No AI-generated content was used without human review.

## Author contributions

X.L. conceived the study, developed the algorithm, performed the analyses, and wrote the manuscript.

## Supplementary Figures

**Supplementary Figure 1.**
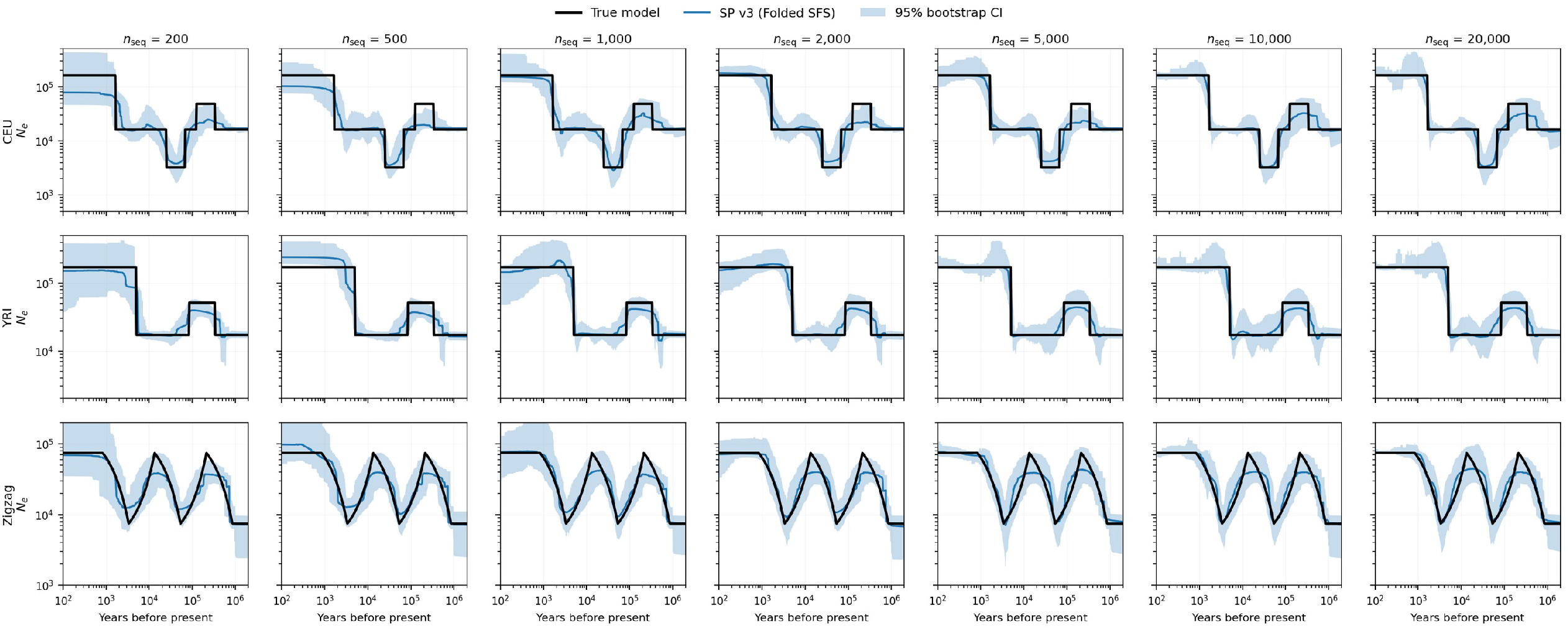
Stairway Plot v3 accuracy across sample sizes using the folded SFS. Inferred *N*_*e*_ (*t*) trajectories (blue, with 95% bootstrap CIs shaded) for three demographic models (rows: CEU, YRI, zigzag) at seven sample sizes (columns: *n*_seq_ = 200 to 20, 000), all using the folded SFS. The true model is shown in black. Accuracy improves and confidence intervals tighten progressively with increasing sample size. At *n*_seq_ *≥* 5, 000, all three models are recovered with high fidelity, including the rapid oscillations of the zigzag model. All simulations used *μ* = 1.2 × 10^−8^, *r* = 1.2 × 10^−8^, *g* = 24 years, and *L* = 300 Mb (10 chromosomes × 30 Mb).

**Supplementary Figure 2.**
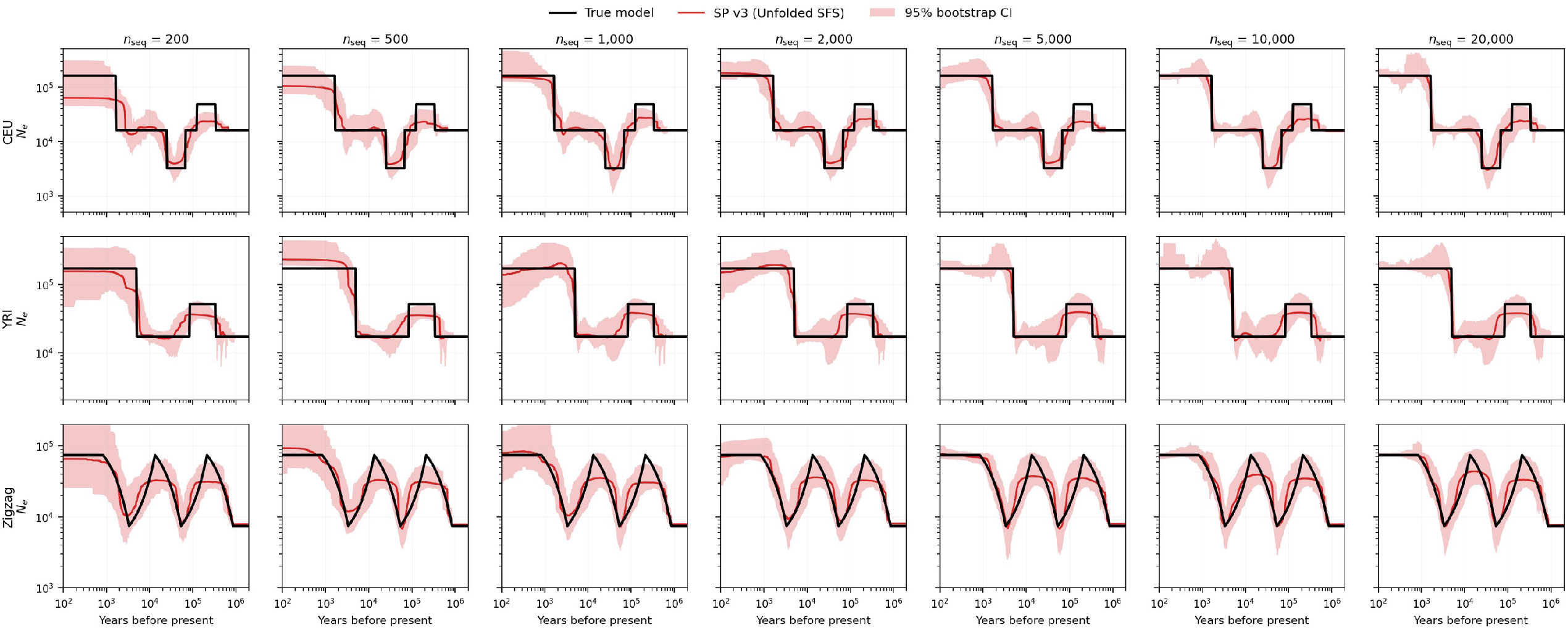
Stairway Plot v3 accuracy across sample sizes using the unfolded SFS. Same layout as Supplementary Figure 1, but using the unfolded SFS (red). Accuracy similarly improves with sample size, but confidence intervals are generally wider than those from the folded SFS (Supplementary Figure 1), particularly at ancient time depths. Because ancestral alleles are known exactly in simulation, folding discards information but does not introduce bias. The narrower folded bands are empirical bootstrap bands and are not interpreted as evidence of greater precision; coverage was not evaluated. For real data, where ancestral-allele assignment is imperfect, we recommend the folded SFS.

## Supplementary Methods: Stairway Plot v3

### Overview of the Stairway Plot Framework

Stairway Plot (Liu & Fu, 2015; Liu & Fu, 2020) infers the history of effective population size *N*_*e*_ (*t*) from the site frequency spectrum (SFS). The SFS summarizes genetic variation in a sample of *n* sequences (haplotypes): for each derived allele count *k* ∈ {1, …, *n* − 1}, the entry *c*_*k*_ records the number of segregating sites at which exactly *k* sequences carry the derived allele.

### Piecewise-constant *N*_*e*_ model

Under the Kingman coalescent (Kingman, 1982), a sample of *n* lineages undergoes *n* − 1 coalescent events as one traces back in time, reducing the lineage count from *n* to *n* − 1 to *…* to 1 (the MRCA). These events divide the time axis into *n* − 1 epochs, where epoch *i* (*i* = *n, n* − 1, …, 2) is the interval during which there are exactly *i* ancestral lineages. Stairway Plot models *N*_*e*_ (*t*) as piecewise-constant with change points at these coalescent event times, a formulation also known as the constant-in-state model (Strimmer & Pybus, 2001; Drummond et al., 2005; Fu, 2025a; Fu, 2025b; Fu, 2026).

To reduce the number of free parameters, contiguous epochs are grouped (Supplementary Methods Figure 1A). Group *j* spans a set of consecutive epochs and assigns them a common scaled mutation rate *θ* _*j*_ = 4 *N*_*e, j*_ *μ*, where *μ* is the per-site per-generation mutation rate. The placement of group boundaries (breakpoints) determines the model complexity: more breakpoints allow finer resolution of *N*_*e*_ changes, but risk overfitting.

### SFS likelihood

The expected proportion of segregating sites at frequency *k* is a linear function of *θ* (Fu, 1995):

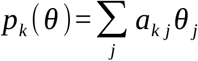

where *a*_*k j*_ is the expected total branch length in the coalescent tree that subtends exactly *k* leaves and falls within the epochs covered by group *j* (Supplementary Methods Figure 1B,C). If group *j* spans the coalescent epochs with *i*∈ {*i*_*st ar t*_,*…, i*_*end*_} lineages, then (Fu, 1995):

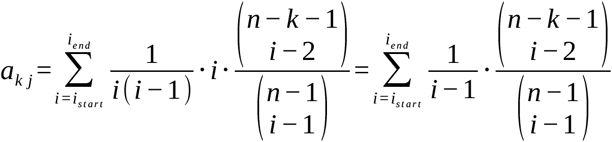

where 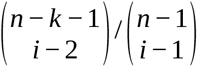 is the probability that a mutation on a branch subtending *i* lineages has derived allele count *k* in the sample, and 1 / [*i* (*i* − 1)] is the expected epoch duration (in units of 4 *N*_*e*_ generations) when there are *i* lineages. The factor of *i* accounts for the *i* branches present during the epoch. Given observed counts *c*_*k*_ from *L* total sites, the log-likelihood is the standard multinomial form:

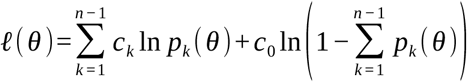

where 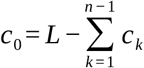 is the number of monomorphic sites. Since *p*_*k*_ is linear in *θ* and ln(⋅) is concave, *l* is concave in *θ* (see Section 1).

Supplementary Methods Figure 1. The Stairway Plot piecewise-constant model. (A) *N*_*e*_ (*t*) is modeled as piecewise-constant with change points at coalescent events (Strimmer & Pybus, 2001; Drummond et al., 2005). Contiguous epochs are grouped; each group shares a common *θ* _*j*_ and is shown in the same color. Time increases upward. (B) The contribution matrix *a*_*k j*_: each SFS bin *k* receives contributions from multiple *θ* groups, with the relative weight depending on the expected branch length at frequency *k* within each group’s coalescent intervals (Fu, 1995). Low-frequency bins (small *k*) are informed by all groups; high-frequency bins are dominated by the most ancient group. (C) The multinomial log-likelihood is linear in *θ* inside the logarithm, making *l* concave—the key property exploited by the convex solver in v3.

### Model selection

Stairway Plot uses a data-driven approach to select the number and placement of breakpoints:

1. **Random candidate breakpoints**. A set of *n*_*rand*_ candidate breakpoint positions is drawn at random from the *n*−2 possible positions.
2. **Training/testing split**. The SFS bins are randomly split into a training set (*p*_*trainin g*_ = 0.67) and a testing set, used to prevent overfitting during model selection.
3. **Forward greedy search**. Starting from a single group (*d i m* = 1), the algorithm iteratively adds the breakpoint that most improves the training-set log-likelihood. At each step, the *θ* parameters are optimized given the current group structure. Forward search stops when the testing-set log-likelihood improvement falls below a threshold (*l r t T hr e s hol d* = 1.0).
4. **Replication**. Steps 1–3 are repeated 200 times with different random breakpoint sets and different training/testing splits. The 200 fitted trajectories are summarized by their median and percentile bands (2.5%, 12.5%, 87.5%, 97.5%) at each time point.

In v2 (Liu & Fu, 2020), this procedure is run at the user’s native sample size *n* with multiple user-specified *n*_*r and*_ values (typically *n* / 4, *n* / 2, 3 *n* / 4, *n* − 2). Candidate breakpoints are drawn uniformly among SFS bin indices. The *θ* optimization uses Differential Evolution (DE; Storn & Price, 1997), a stochastic metaheuristic. An autocorrelation constraint limits the ratio between adjacent *θ* values to prevent extreme jumps.

### Changes in v3

v3 introduces five categories of improvements over v2. The changes span: (1) a convex interior-point solver replacing differential evolution, (2) breakpoint sampling in coalescent-time space, (3) multi-projection sigmoid stitching, (4) per-replicate SFS bootstrap with random hypergeometric projection, and (5) backward–swap model refinement with smoothness regularization.

## 1. Convex Interior-Point Solver

In v2, DE treats the log-likelihood as a black-box objective, does not exploit the mathematical structure of the problem, is not guaranteed to find the global optimum, and can be slow as the number of *θ* parameters grows. v3 replaces DE with an exact solver that exploits the concavity of the SFS log-likelihood.

### Concavity of the SFS log-likelihood

As shown in the SFS likelihood section above, 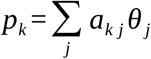is linear in *θ*. The log-likelihood is therefore a sum of terms 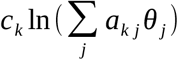. Since each *c*_*k*_ *≥* 0, each *a*_*k j*_ *≥* 0, and ln(⋅) is concave, and a non-negative weighted sum of concave functions composed with affine functions is concave, the entire log-likelihood *l* (*θ*) is concave in *θ*. This guarantees that any local maximum is the global maximum.

### Interior-point solver

Exploiting this concavity, v3 replaces DE with a Newton-based interior-point (barrier) method (Boyd & Vandenberghe, 2004, Ch. 11). The barrier objective is:

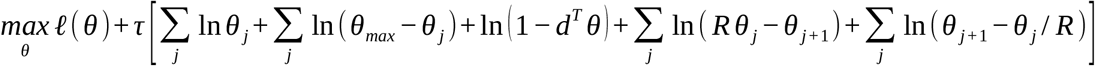

where *l* (*θ*) is the SFS log-likelihood, *τ* > 0 is the barrier parameter, *d* is the vector of constraint coefficients 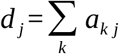, the total contribution of *θ* group *j* across all SFS bins), and *R* = − ln [(1 *− p*_*ac*_) / 2] with *p*_*ac*_ = 0.99 (giving *R ≈* 5.3) is the maximum allowed ratio between adjacent *θ* values (the autocorrelation constraint, see Section 5). The log terms enforce: (i) positivity *θ* _*j*_ > 0, (ii) upper bound *θ* _*j*_ < *θ*_*max*_, (iii) the linear constraint *d*^*T*^ *θ* < 1 (the total expected proportion of segregating sites must be less than 1, ensuring the monomorphic class retains positive probability), and (iv) the autocorrelation constraints *θ* _*j*_ / *R ≤θ*_*j* + 1_ *≤ Rθ* _*j*_. Since each log-barrier term is concave in *θ*, the entire barrier objective remains concave and has a unique maximum for any *τ* > 0 (Boyd & Vandenberghe, 2004, Section 11.2). The solver proceeds in two nested loops: the outer loop decreases *τ* across iterations (e.g., *τ* ←*τ* / 10), gradually thinning the barriers so the solution approaches the true constrained optimum; the inner loop, at each fixed *τ*, maximizes the barrier objective via Newton’s method using the exact gradient and Hessian of the log-likelihood.

The convex solver provides three advantages over DE: (i) guaranteed global optimality for the fixed-grouping subproblem (model selection over breakpoint placements remains heuristic and is explored by replicated randomized searches), (ii) faster convergence (typically 10–20 Newton iterations vs. thousands of DE evaluations), and (iii) exact gradient/Hessian information enables the smoothness and curvature penalties described below.

## 2. Log-Coalescent-Time Breakpoint Sampling

In v2, candidate breakpoints are drawn uniformly among bin indices {3,*…, n*_*seq*_}, giving each frequency class equal probability. However, coalescent time depth increases rapidly with decreasing frequency class *k*. We define *t* (*k*) as the expected time (in years) for a sample of *n* lineages to coalesce down to *k* lineages under constant *N*_*e*_ (Kingman, 1982):

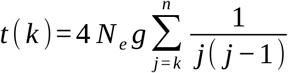

where *g* is the generation time (years per generation). Consequently, uniform sampling in bin-index space produces far more breakpoints in recent time (high *k*, short *t*) than in ancient time (low *k*, long *t*), leading to over-parameterization of recent epochs and under-resolution of ancient ones (Supplementary Methods Figure 2A).

Supplementary Methods Figure 2. Log-coalescent-time breakpoint sampling. (A) v2 uniform sampling in bin index: breakpoints (red ticks) cluster in the middle log (*t*) range where SFS bins are densest, leaving ancient times with zero breakpoints. Gray ticks show all available SFS bin positions. Numbers above show breakpoint counts per 0.5 units of log_10_ (*t*). (B) v3 uniform sampling in log (*t*): breakpoints (green ticks) are spread evenly across the full time range, with comparable counts per interval. (C) Side-by-side density comparison confirms that v3 achieves approximately uniform coverage while v2 concentrates breakpoints in a narrow band. (D) The recommended *n*_*r and*_ = *max* (20, ⌊ 5 log_2_ *n*_*s e q*_ ⌉) (blue) tracks the log-time window width *W* (*n*)= 2 log_10_ *n* (orange), maintaining a constant density of ~8.3 breakpoints per unit of log_10_ (*t*) as sample size grows.

### Change in v3

We sample candidate breakpoints uniformly in log_10_ *t* (*k*) space. Specifically:

1. Compute log_10_ *t* (*k*) for each bin *k ∈* {3,*…, n*_*s e q*_} using the formula above (the constant 4 *N*_*e*_ *g* shifts all values by the same additive constant in log-space and thus does not affect the sampling).
2. Define bounds *L*_*min*_ = log_10_ *t* (*n*_*s e q*_) (most recent) and *L*_*max*_ = log_10_ *t* (3) (most ancient).
3. Draw a target uniformly: *u* ~ U n*i f or m* [*L*_*min*_, *L*_*max*_].
4. Map to the nearest SFS bin: *k*\*= *arg min*_*k*_ | log_10_ *t* (*k*)*− u* |.
5. Repeat until *n*_*r and*_ unique breakpoints are selected.

Because the targets *u* are drawn uniformly in log_10_ (*t*), the resulting breakpoints have approximately equal density per unit of log_10_ (*t*) (Supplementary Methods Figure 2B,C), ensuring that the model has the resolution to detect demographic events at any time depth.

### Recommended *n*_*r and*_

We replace the user-specified *n*_*r and*_ values of v2 (typically *n* / 4, *n* / 2, 3 *n* / 4, *n* − 2) with a single data-driven formula:

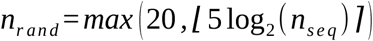

This yields *n*_*r and*_ = 20 at *n*_*se q*_ = 16, 33 at *n*_*se q*_ = 100, 50 at *n*_*se q*_ = 1000, and 66 at *n*_*se q*_ = 10, 000. The logarithmic scaling ensures approximately constant breakpoint density per unit of log_10_ (*t*) as sample size grows (Supplementary Methods Figure 2D).

## 3. Multi-Projection Sigmoid Stitching

In v2, inference is run at the user’s native sample size *n* with multiple *n*_*r and*_ values and the results are averaged. This provides variance reduction but does not address a fundamental limitation: a single SFS projection at sample size *n* has a finite “trust region” in time (Supplementary Methods Figure 3A,B). It resolves *N*_*e*_ accurately only for epochs whose coalescent timescale matches the sampling depth of that projection. Since all v2 runs share the same sample size, they all share the same trust region.

Supplementary Methods Figure 3. Motivation for multi-projection sigmoid stitching. (A) SFS bins map nonlinearly to coalescent time: the relationship between bin index *k* and time depth *t* (*k*) is steep, with high-*k* bins compressed into recent time and low-*k* bins spanning ancient time. (B) Positions of individual SFS bins on the log_10_ (*t*) axis for three projection sizes. Bins are sparse at both ends and densest in the middle range (brackets show the middle 50% of bins, where resolution is highest). Each projection’s dense region covers a different time interval, motivating the use of multiple projections. (C) Geometric projection spacing produces even coverage in log (*t*); linear spacing crowds projections at the recent end, leaving ancient time under-resolved. (D) Sigmoid stitching weights for a geometric projection set: each projection dominates its trust region with smooth handoffs.

v3 addresses this by running inference independently at multiple *projection sizes* (subsamples of the original SFS), then combining the per-projection *N*_*e*_ (*t*) trajectories using a sigmoid-weighted average in log-time space.

### Default projection set

Given a user SFS at sample size *n*, we use the geometric set {20, 50, 100, 200, 500, 1000, 2000, 5000, 10, 000, 20, 000, 50, 000, 100, 000, 200, 000} capped at *n*. Small projections resolve ancient time; large projections resolve recent time.

The projections are geometrically spaced (approximately constant ratio between consecutive values) rather than linearly spaced, ensuring that each projection’s trust region has approximately equal width in log (*t*) (Supplementary Methods Figure 3C,D).

### Trust region boundaries

For *P* projections sorted largest-to-smallest as *n*_1_ > *n*_2_ > > *n*_*P*_, the trust region 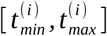 for projection *n*_*i*_ is:

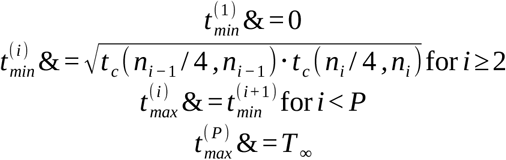

Where 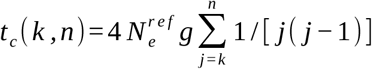 is the coalescent time to reach *k* lineages from *n*, 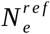 is the median inferred *N*_*e*_ from the largest projection (or user-specified), and *T*_*∞*_ = 10^8^ years.

### Sigmoid weighting

At each time point on a log-uniform grid, the weight for projection *n*_*i*_ is:

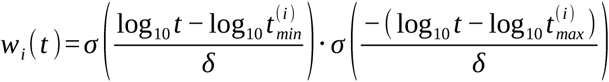

where *σ* (*x*)= 1 / (1 + *e*^*− x*^) is the logistic function. The transition width *δ* is set to the median log-width of interior projections:

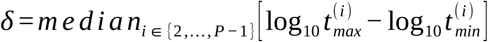

Weights are normalized to sum to 1 at each time point.

### Geometric average

The stitched *N*_*e*_ (*t*) trajectory and its confidence intervals are computed as the weighted geometric (log-space) mean:

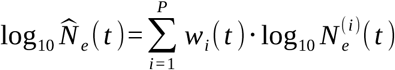

where 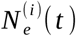 is the inferred *N*_*e*_ at time *t* from projection *n*_*i*_, interpolated onto the common log-uniform time grid.

## 4. Per-Replicate SFS Bootstrap and Random Projection

In v2, all 200 replicate fits share the same input SFS. The inter-replicate variance arises solely from the random choice of breakpoint positions and the random training/testing split. This captures *fitting variance* (sensitivity to the model structure) but misses *sampling variance* (multinomial resampling variability of the frequency-class counts, conditional on the observed number of segregating sites) and *projection variance* (stochasticity from subsampling *n*^*′*^ from *n* haplotypes).

In v3, each of the 200 replicates at each projection receives its own stochastic SFS, generated by a two-step pipeline:

### Step 1: Multinomial bootstrap

Given the observed SFS *c* =(*c*_1_, …, *c*_*n*−1_) at the native sample size *n* with 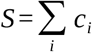 total segregating sites, draw:

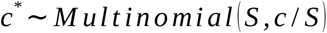

This is the standard nonparametric bootstrap (Efron, 1979) applied to the SFS: resampling *S* segregating sites with replacement from the empirical distribution over frequency classes. The total number of segregating sites S is held at its observed value, so the procedure conditions on S; the monomorphic count enters the likelihood through L but is not resampled. Sites are treated as independent, so the resampling does not capture linkage among sites.

### Step 2: Random hypergeometric projection

For each frequency class *k* in the bootstrapped SFS *c*\* at sample size *n*, and target projection size *n*^*′*^:

- For each of the 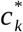 sites in bin *k*, draw the projected count *k*^*′*^ ~ *Hypergeometric* (*n, k, n*^*′*^).
- Accumulate counts into the target SFS: if 1 *≤ k*^*′*^ *≤n*^*′*^ − 1, add to *S F S*_*n*_^*′*^ [*k*_*′*_]; otherwise the site becomes monomorphic in the subsample and is excluded.

This replaces the deterministic expectation-based projection of earlier versions, where 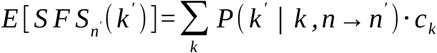 produced identical projected SFS across all replicates.

### Seed derivation

To ensure bit-for-bit reproducibility regardless of the execution order of replicates (critical for cluster-array dispatch), each (*s, n*^*′*^, *r*) triple maps to a unique deterministic seed pair via a hash:

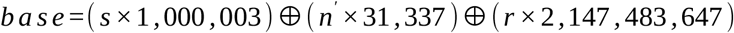

where ⊕ is bitwise XOR. The base initializes a PCG64 generator that produces the Java and Python seed pair for that replicate.

### Effect on confidence intervals

The 200 replicates at each projection now vary due to three sources:

- SFS sampling variance (multinomial bootstrap),
- Projection variance (hypergeometric draw),
- Fitting variance (random breakpoints + training/testing split).

In v2, only the third source contributed. In v3, the first two sources are added, so the CIs additionally reflect multinomial resampling of the frequency-class counts (conditional on the observed number of segregating sites and assuming independent sites) and stochastic subsampling. They do not capture linkage among sites or uncertainty in the total number of segregating sites; a genomic block bootstrap would be required for that.

### CI computation

At each projection *n*_*i*_, the 200 replicate fits each produce a piecewise-constant *N*_*e*_ (*t*) trajectory. These are aggregated by the Java summary stage (Stairway_output_summary_plot2), which sorts the 200 *N*_*e*_ values at each time point and reports the median, 2.5th, 97.5th, 12.5th, and 87.5th percentiles as the per-projection .final.summary file. During stitching, each percentile column is interpolated onto the common log-uniform time grid and combined independently using the same sigmoid weights:

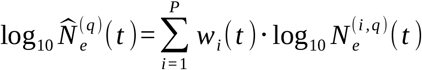

where *q* ∈ {2.5 %, 12.5 %, 50 %, 87.5 %, 97.5 %} and 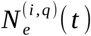 is the *q*-th percentile from projection *n*_*i*_. The stitched 2.5th–97.5th percentile band is reported as the 95% CI, and the 12.5th–87.5th band as the 75% CI.

## 5. Model Selection and Regularization

The forward greedy search, training/testing split, and autocorrelation constraint described in the Overview are shared between v2 and v3. The autocorrelation constraint limits adjacent *θ* ratios to a factor *R*:

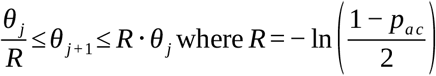

with *p*_*ac*_ = 0.99 by default, giving *R ≈* 5.3. v3 adds the following refinements, which are independent of the solver choice but become practical with the convex solver (Section 1) because each candidate configuration can be re-optimized in milliseconds rather than minutes:

### Backward elimination

After forward search terminates, iteratively attempt removing each split position. Accept removal if neither the training-set nor testing-set fitness worsens by more than *l r t T hr e s hol d*. This eliminates spurious breakpoints that were beneficial at an earlier model dimension but become redundant once neighboring splits are in place.

### Swap moves

After backward elimination, try relocating each remaining split to a nearby allowed position (within its adjacent boundaries). Accept the move if it improves training fitness and the testing-set improvement exceeds *lrtThreshold*. This fine-tunes breakpoint placement without changing model dimension.

### Warm-starting

When testing a candidate split at position *j* that divides group *g* into two sub-groups, the solver is initialized with 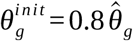 and 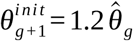, where 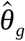 is the current best estimate for the parent group. This accelerates convergence by starting near the expected solution.

### Smoothness penalty

The convex solver incorporates a quadratic first-difference penalty:

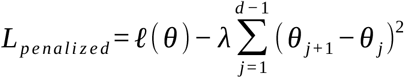

where *l* (*θ*) is the SFS log-likelihood and *λ* is the penalty strength. This discourages abrupt jumps between adjacent *θ* parameters, suppressing high-frequency oscillations (“ringing”) at sharp demographic transitions. The penalty is applied in *θ*-space rather than log *θ*-space, which preserves the concavity of the penalized objective (a quadratic penalty on log-transformed variables would not be concave). Because *θ*∝ *N*_*e*_, this penalizes absolute rather than relative changes; the autocorrelation constraint (ratio bound *R*) serves as a complementary scale-invariant control.

### Auto-tuned *λ*

Rather than fixing *λ*, the value is selected adaptively from a grid {0, 10^4^, 10^5^, 10^6^, 10^7^} at each dimension increase. The winning candidate is resolved at each grid value, and the *λ* minimizing the testing-set fitness is adopted. The log-likelihood scale changes with the number of segregating sites, but because both the training-set objective and the testing-set selection criterion scale with data size, the grid remains effective across the range of simulated and real datasets used here.

### Curvature penalty

An additional second-difference (curvature) penalty is applied at *λ*_*cur v*_ = *λ* / 10:

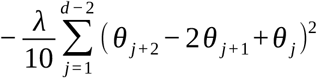

While the smoothness penalty discourages jumps between adjacent *θ* values, the curvature penalty additionally discourages oscillations—rapid alternation between high and low values. The curvature penalty is set to 1/10 of *λ* so that it acts as a secondary correction.

## 6. Summary of Changes

| Component | v2 | v3 |
| --- | --- | --- |
| Breakpoint sampling | Uniform in bin index | Uniform in $\log t_c$ |
| $n_{rand}$ | User-specified<br>(e.g. $n/4, n/2, \dots$) | $\max(20, \lfloor 5 \log_2 n_{seq} \rfloor)$ |
| Projections | Single (native $n$) | Multi-projection<br>geometric set |
| Combination | Average over $n_{rand}$<br>values | Sigmoid stitching in<br>$\log t$ |
| SFS per replicate | Identical<br>(deterministic) | Bootstrapped +<br>random projection |
| CI sources | Fitting variance only | Conditional SFS<br>resampling +<br>projection + fitting |
| Optimizer | Differential<br>Evolution<br>(stochastic) | Interior-point<br>(convex; global<br>optimum for fixed<br>grouping) |
| Model search | Forward only | Forward + backward<br>+ swap |
| Regularization | Autocorrelation<br>constraint only | Autocorrelation +<br>smoothness +<br>curvature, auto-λ |
| Solver warm-start | No | Split-aware warm-start |

### Implementation

All changes are implemented in the following source files of the Stairway Plot v3 distribution (https://github.com/xiaoming-liu/stairway-plot-v3):

- stairway_plot_es/Stairway_fold_training_testing9_logtime.java and Stairway_unfold_training_testing9_logtime.java: Core inference engine with log-time breakpoint sampling, forward–backward–swap search, smoothness/curvature penalties, auto-*λ*, and warm-starting.
- stairway_plot_es/ConvexThetaSolver.java: Interior-point barrier solver with smoothness and curvature penalty terms.
- lib/sfs_project.py: Multinomial bootstrap and random hypergeometric projection.
- lib/stitch.py: Multi-projection sigmoid stitching with trust-region computation.
- lib/pipeline.py: Orchestration of per-replicate seed derivation, SFS preparation, Java dispatch, and post-processing.
- stairway_plot_v3.8.3.py: Command-line entry point with subcommands (run, prepare-cluster, collect, etc.).

